## Extended Data Figures and Supplementary Table legends for "RNA language models predict mutations that improve RNA function"

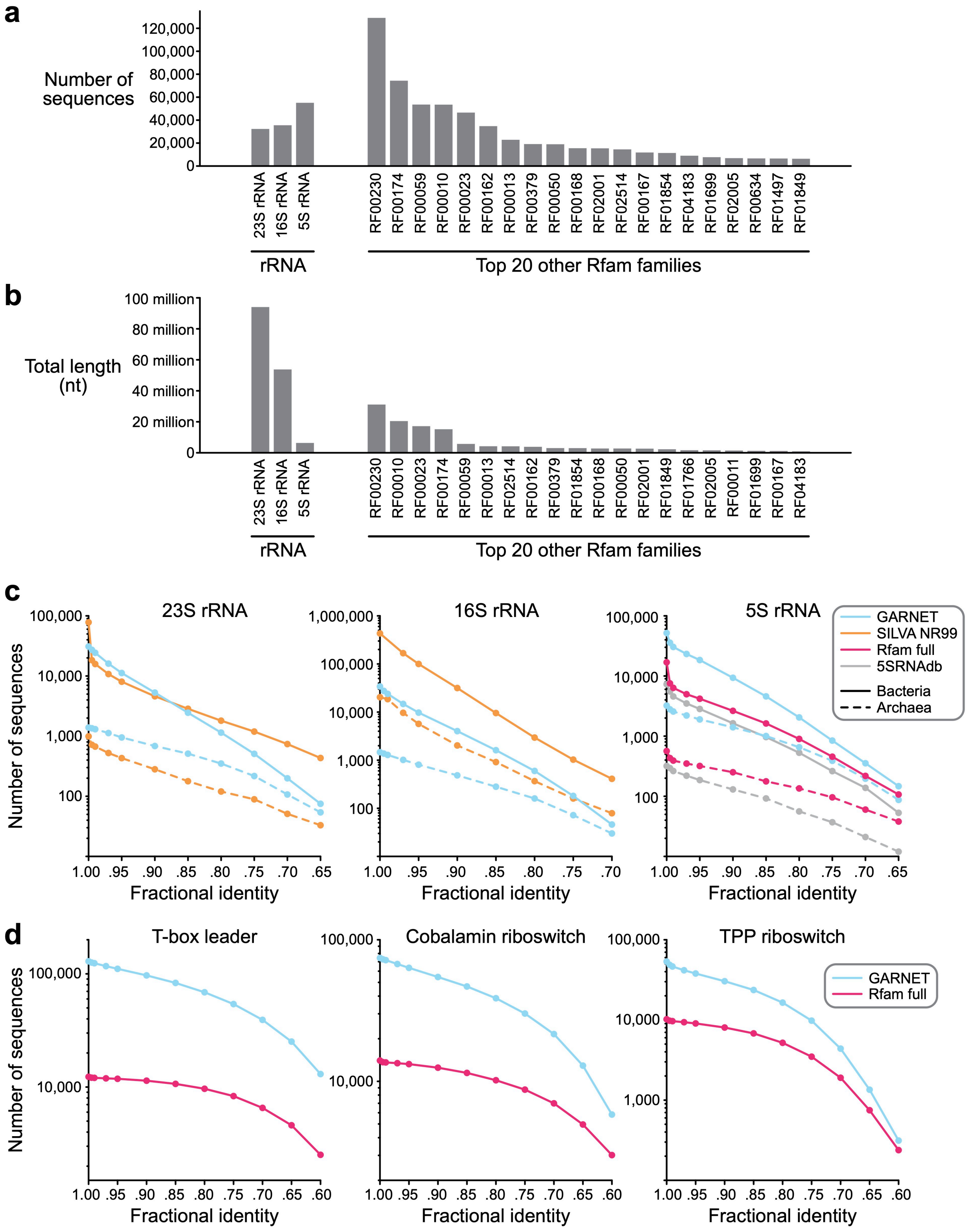

**Extended Data Fig. 1: Additional evaluation of RNA dataset sequence diversity in GARNET. a.** Number of GARNET sequences for rRNA and for the top twenty most abundant of the 228 RNA families. **b.** Total sequence length of GARNET RNA sequences for rRNA and for the top twenty most abundant of the 228 RNA families. **c.** Comparing diversity of GARNET-based alignments against state-of-the-art alignments for 23S rRNA, 16S rRNA, 5S rRNA by filtering the alignments at a range of pairwise fractional identity thresholds with esl-weight, part of the HMMER suite of programs[57]. **d.** Diversity comparison for the three most abundant of the 228 RNA families in GARNET with esl-weight.

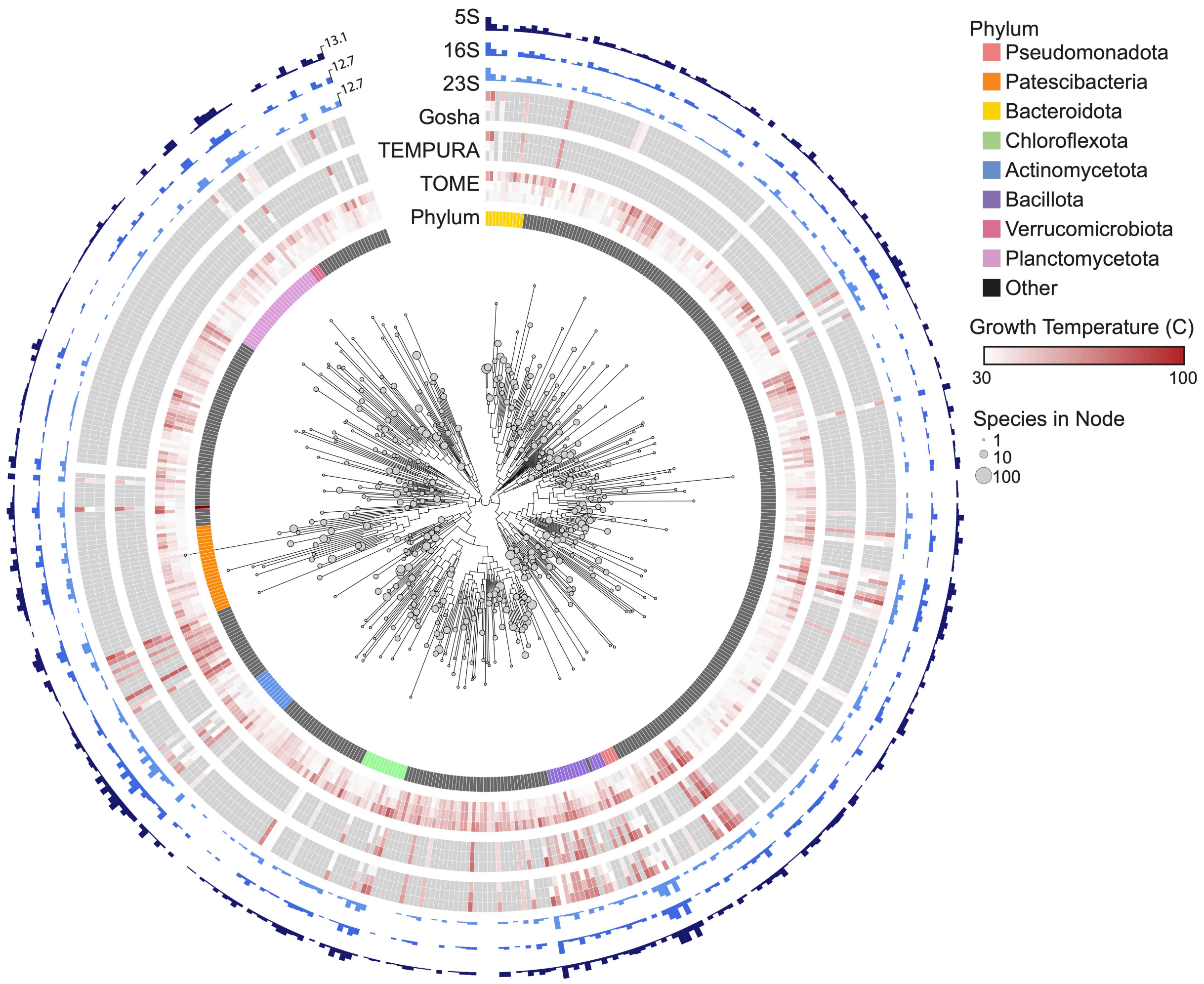


**Extended Data Fig. 2: Bacterial phylogeny within the GTDB including OGTs.** Bacterial phylogenetic tree of GTDB reference organisms, grouped at the Class taxonomic rank, arbitrarily rooted. Node tip sizes are proportional to the number of species represented by node ($log_2$ transformed). Inner circle indicates Phylum. The next circle represents TOME-predicted min, median, and maximal optimal growth temperatures of all species within rank. The next two circles similarly represent empirically measured optimal growth temperatures pulled from the Tempura and Gosha datasets, respectively. Outer circles represent the total number of 23S, 16S, and 5S detected in each rank, respectively ($log_2$ transformed).

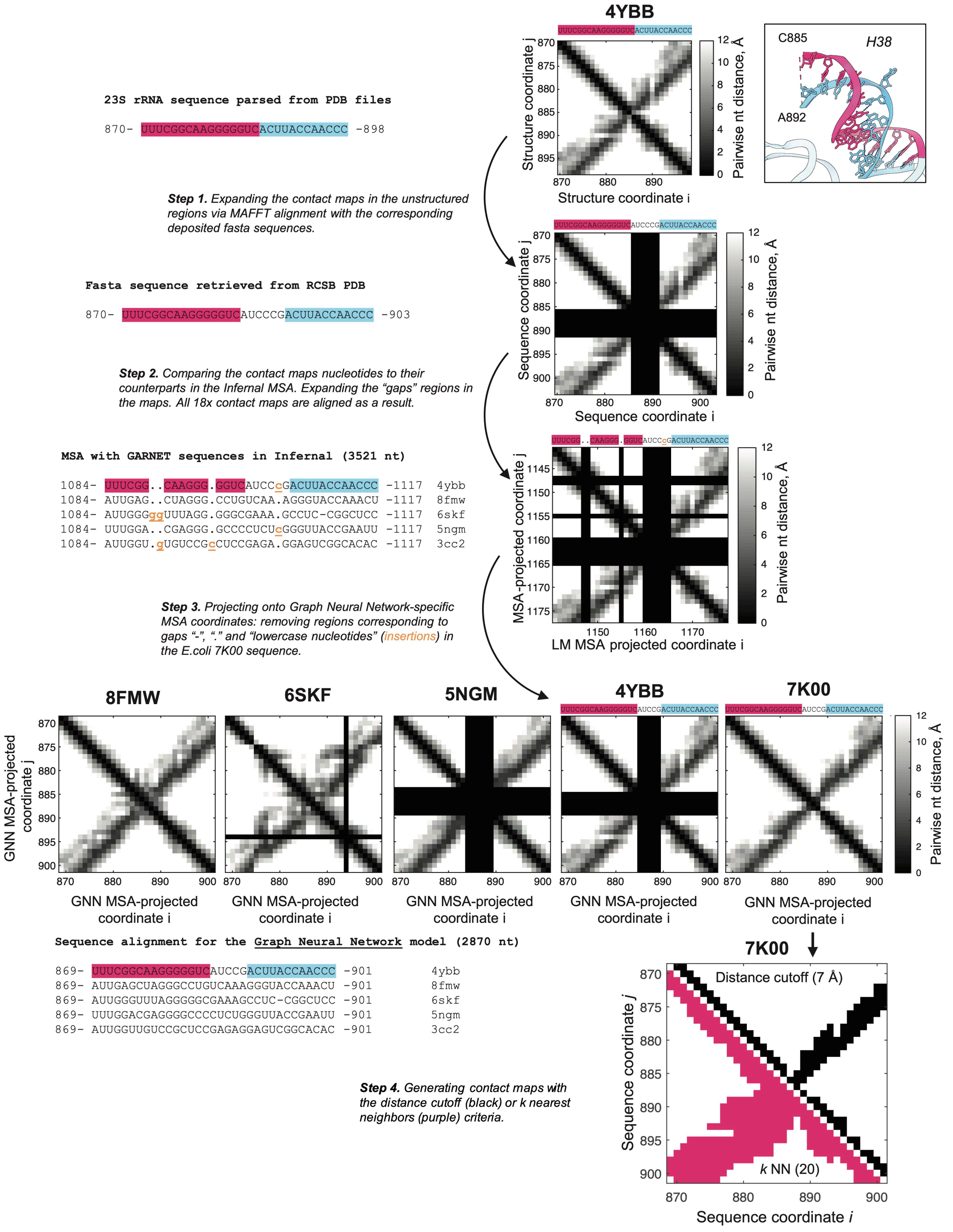


**Extended Data Fig. 3: Schematic of MSA-informed alignment of distance matrices from ribosomal structures and generation of contact maps.** The initial unaligned pairwise nucleotide distance matrices were generated from 23S rRNA structures in the

PDB files (see **Materials and Methods**). Further, sequences of 23S rRNA were extracted directly from the PDB files and corresponding FASTA files available in the Protein Data Bank[38]. In the initial step, nucleotides missing in the structures were pinpointed through a MAFFT alignment comparing FASTA and PDB-derived sequences, leading to the insertion of empty columns and rows at these positions in the distance matrices. Subsequently, in step 2, the extracted archaeal and bacterial rRNA sequences from the structures (shown) were combined with those from GARNET using Infernal, matching the distance matrices' coordinates with the MSA with further introduction of empty rows and columns. At step 3, insertions (lowercase characters in the MSA) and deletions (gaps) were identified in the *E.coli* 7K00 sequence in the MSA, and the corresponding rows and columns were removed from the distance matrices. This process aligned the nucleotides in the distance matrices of all 18 archaeal and bacterial 23S rRNAs with their counterparts in the GARNET-anchored MSA utilized for the GNN model. In the final step, contact maps were generated from the distance matrices based either on the distance cutoff or *k*-nearest neighbors criteria (see **Materials and Methods**).

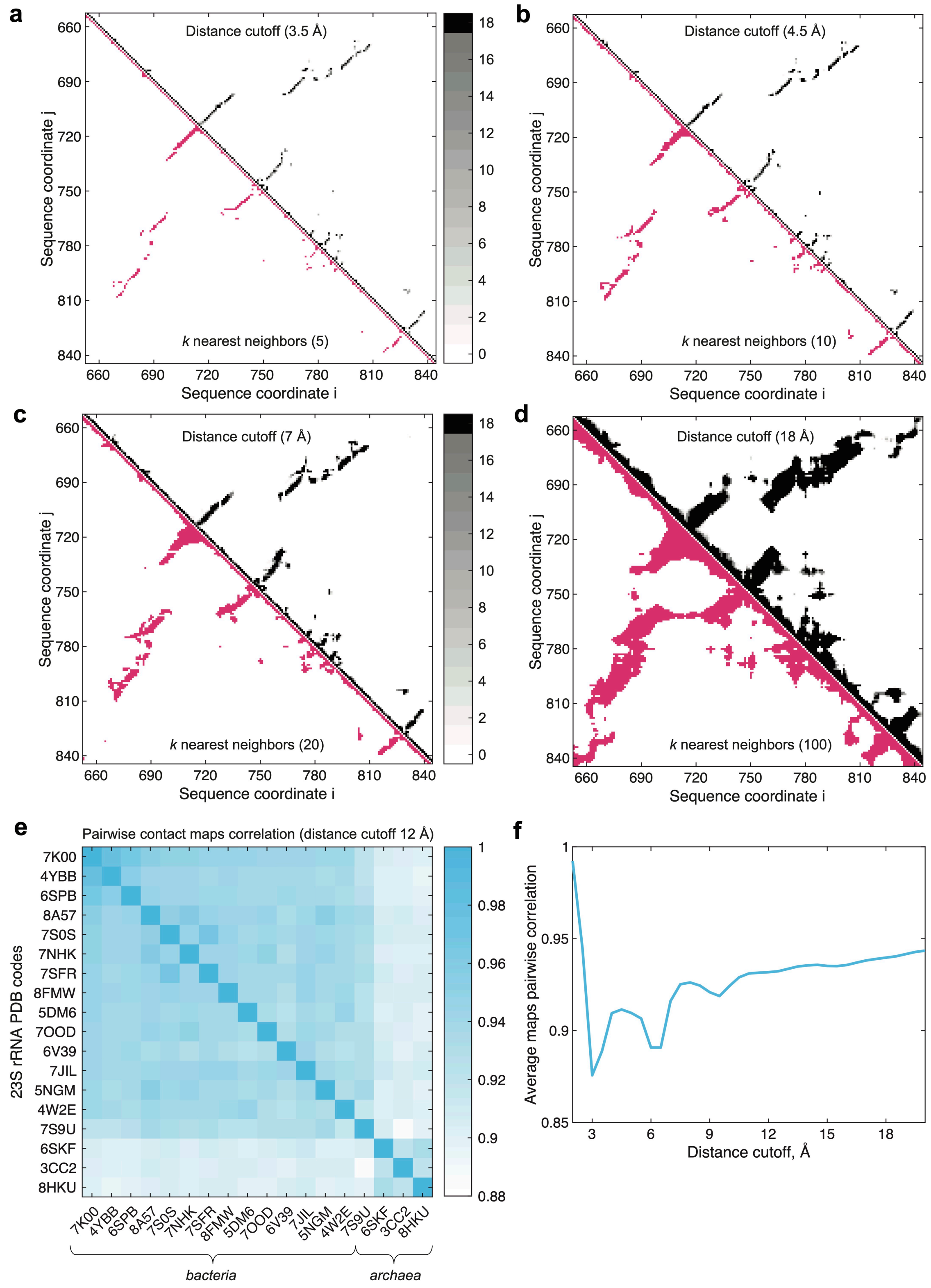


**Extended Data Fig. 4: Contact maps and comparisons of high-resolution bacterial and archaeal 50S subunit structures. a-d.** Comparison of contact maps generated for

23S rRNA with the distance cutoff (top-right) and the *k*-nearest neighbors criteria (bottom-left). The matching pairs of fully sampled distances and *k* values for nearest neighbors were chosen according to the histogram in **Fig. 3d**. **e**. Pairwise correlation of contact maps for 18 bacterial and archaeal 23S rRNA structures, generated at a distance cutoff of 12 Å (see **Materials and Methods**). Note the high degree of structural correlation in the plot, which is also evident from matching of the 18 contact maps feature coordinates in panels (a) through (d). **f.** Average correlation of the 18 contact maps as a function of distance cutoff. Local maxima of correlation at 4.5 Å, 8 Å, 11 Å in the plot correspond to the minima between the peaks in **Fig. 3d**, indicating full sampling of 3.5 Å, 6 Å, 12 Å characteristic internucleotide distances. The structural correlation degree does not rise significantly above the distance cutoff of 12 Å.

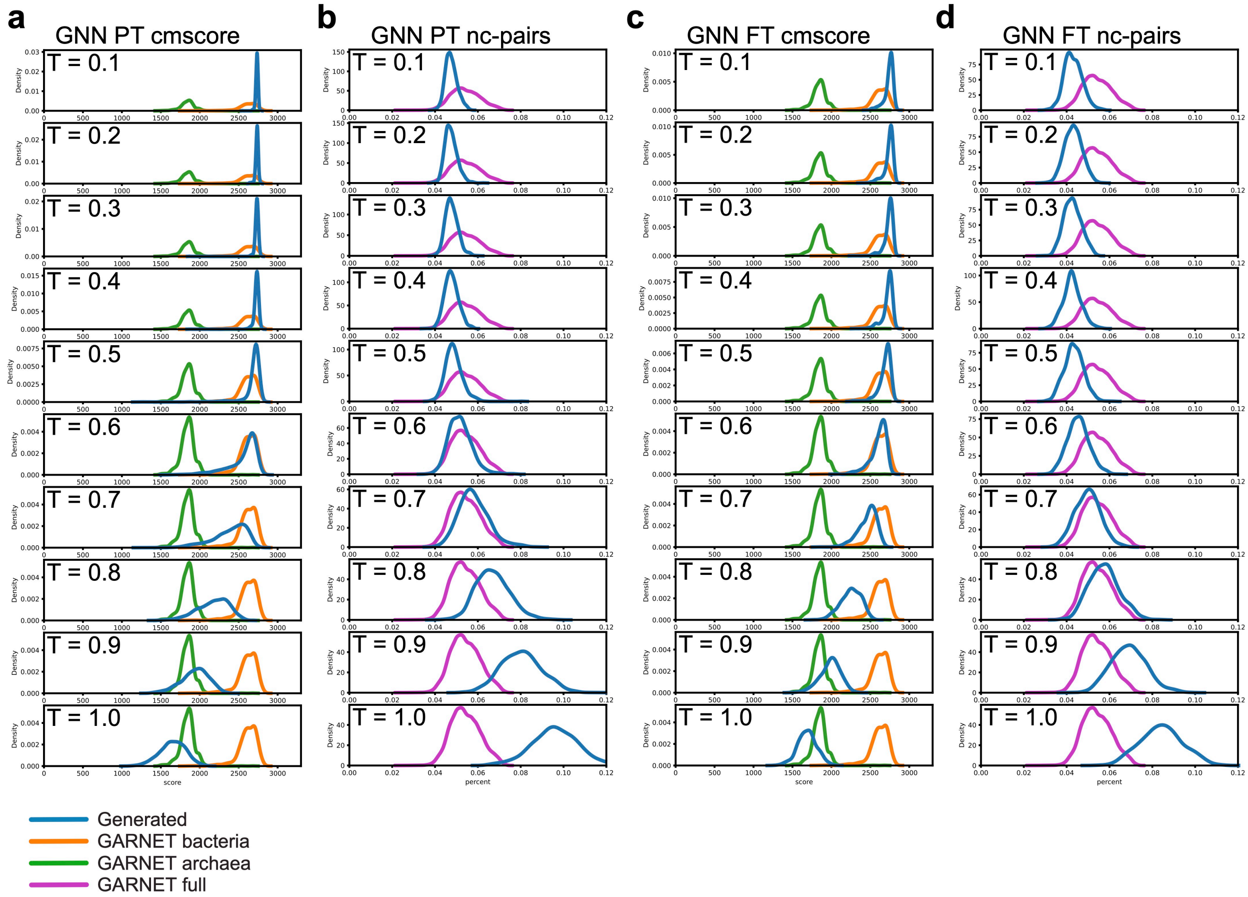


**Extended Data Fig. 5: Properties of RNA sequences generated from the 23S rRNA GNN model. a.** Cmsearch scores for sequences generated from the pretrained GNN model at temperatures ranging from 0.1 to 1.0. **b.** Fraction of mispaired nucleotides of sequences generated from the pretrained GNN model relative to RF02541 at temperatures ranging from 0.1 to 1.0. **c.** Cmsearch scores for sequences generated from the finetuned GNN model at temperatures ranging from 0.1 to 1.0. **d.** Fraction of mispaired nucleotides of sequences generated from the finetuned GNN model relative to RF02541 at temperatures ranging from 0.1 to 1.0.

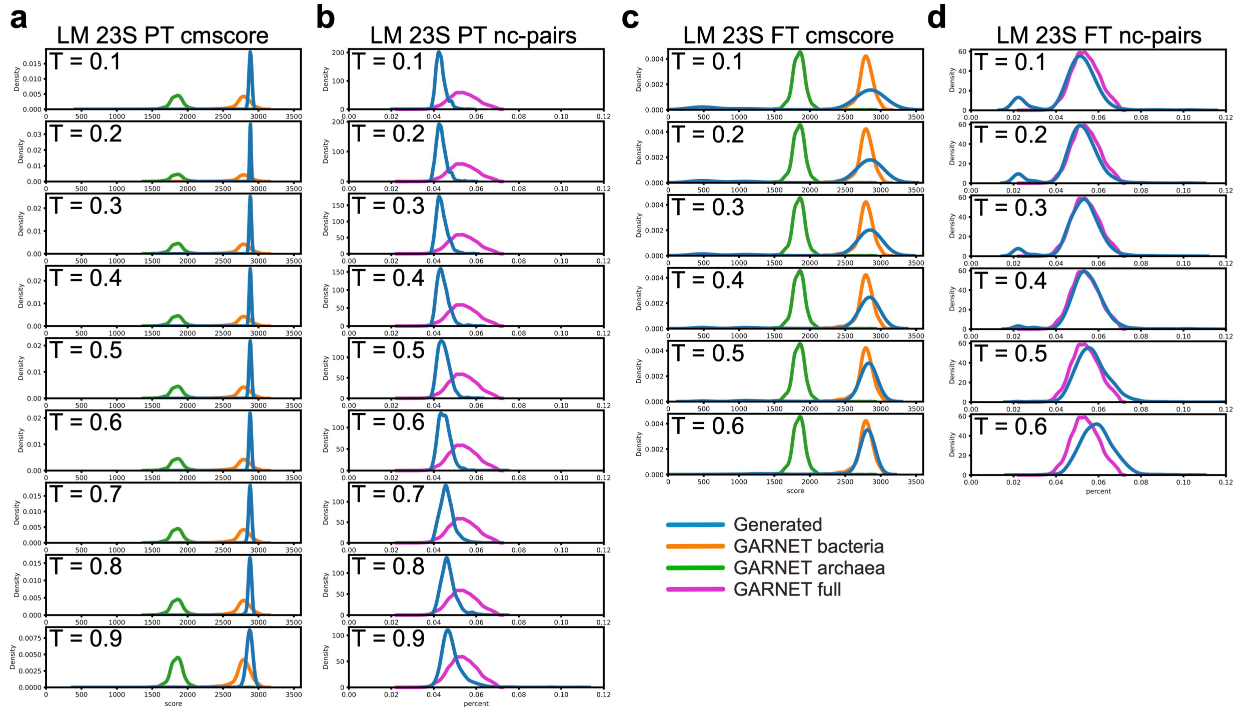


**Extended Data Fig. 6: Properties of RNA sequences generated from the 23S rRNA LM.** **a.** CM scores for sequences generated from the pretrained 23S rRNA LM at temperatures ranging from 0.1 to 0.9. **b.** Fraction of mispaired nucleotides of sequences generated from the pretrained 23S rRNA LM relative to RF02541 at temperatures ranging from 0.1 to 0.9. **c.** CM scores for sequences generated from the finetuned 23S rRNA LM at temperatures ranging from 0.1 to 0.6. **d.** Fraction of mispaired nucleotides of sequences generated from the finetuned 23S rRNA LM relative to RF02541 at temperatures ranging from 0.1 to 0.6.

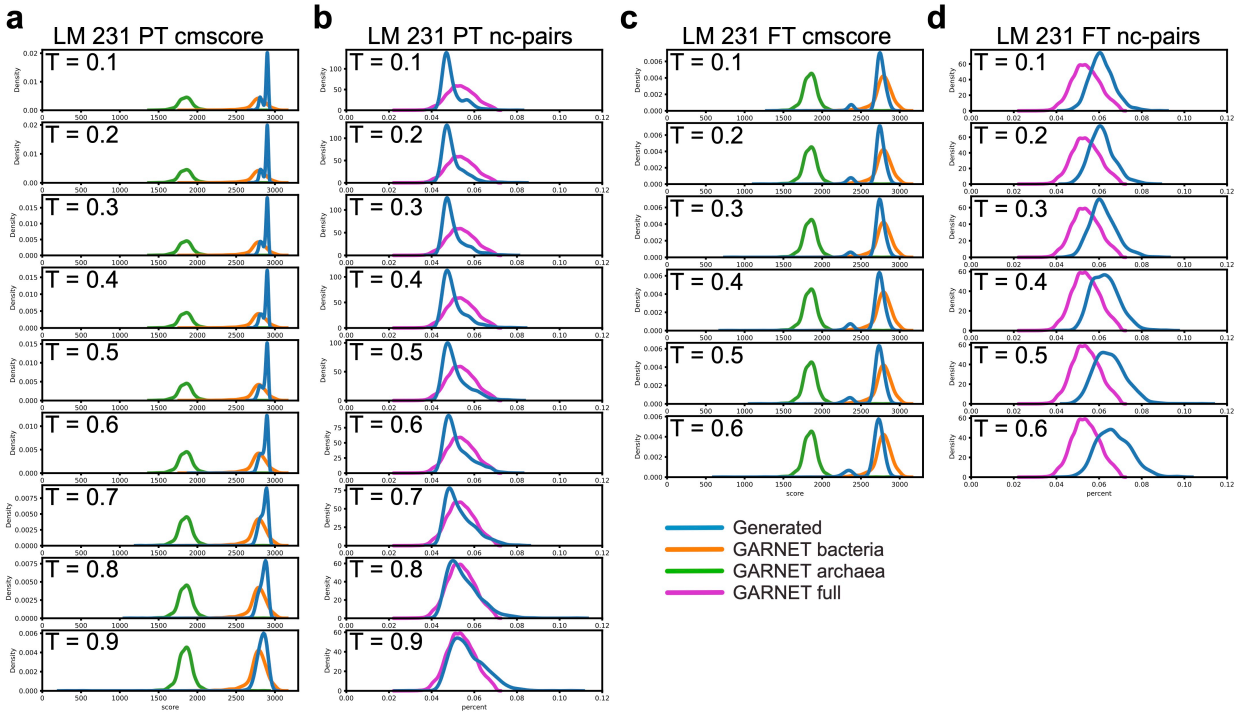


**Extended Data Fig. 7: Properties of RNA sequences generated from the 23S rRNA LM models trained on the 231-RNA dataset. a.** CM scores for sequences generated from the pretrained RNA LM model at temperatures ranging from 0.1 to 0.9. **b.** Fraction of mispaired nucleotides of sequences generated from the pretrained RNA LM relative to RF02541 at temperatures ranging from 0.1 to 0.9. **c.** CM scores for sequences generated from the finetuned RNA LM model at temperatures ranging from 0.1 to 0.6. **d.** Fraction of mispaired nucleotides of sequences generated from the finetuned RNA LM model relative to RF02541 at temperatures ranging from 0.1 to 0.6.

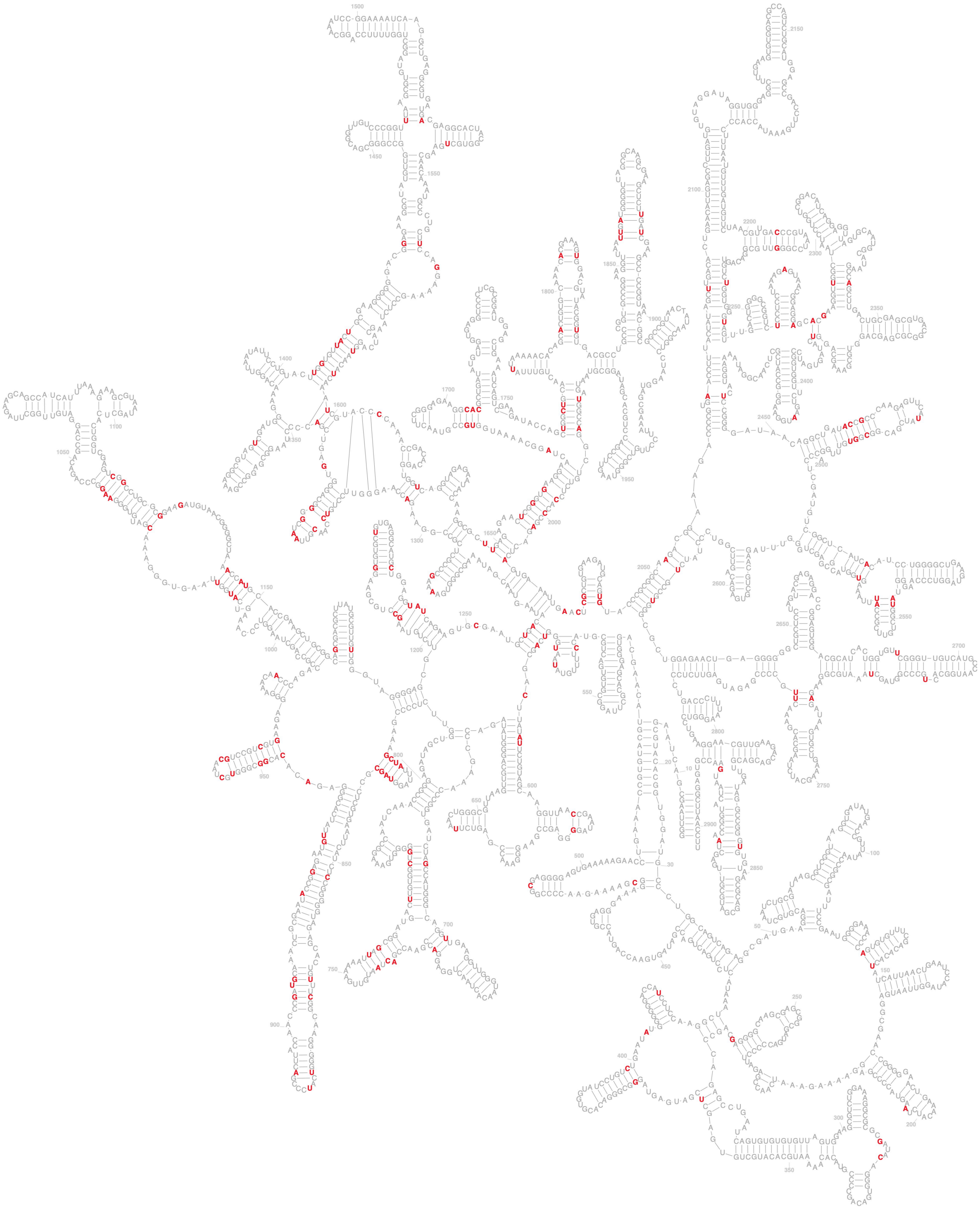

**Extended Data Fig. 8: Top 200 JSD locations from the GNN model.** Top ranking 200 Jensen-Shannon divergence values calculated from a secondary structure alignment of generated *E. coli* 23S rRNA sequences are colored in red.

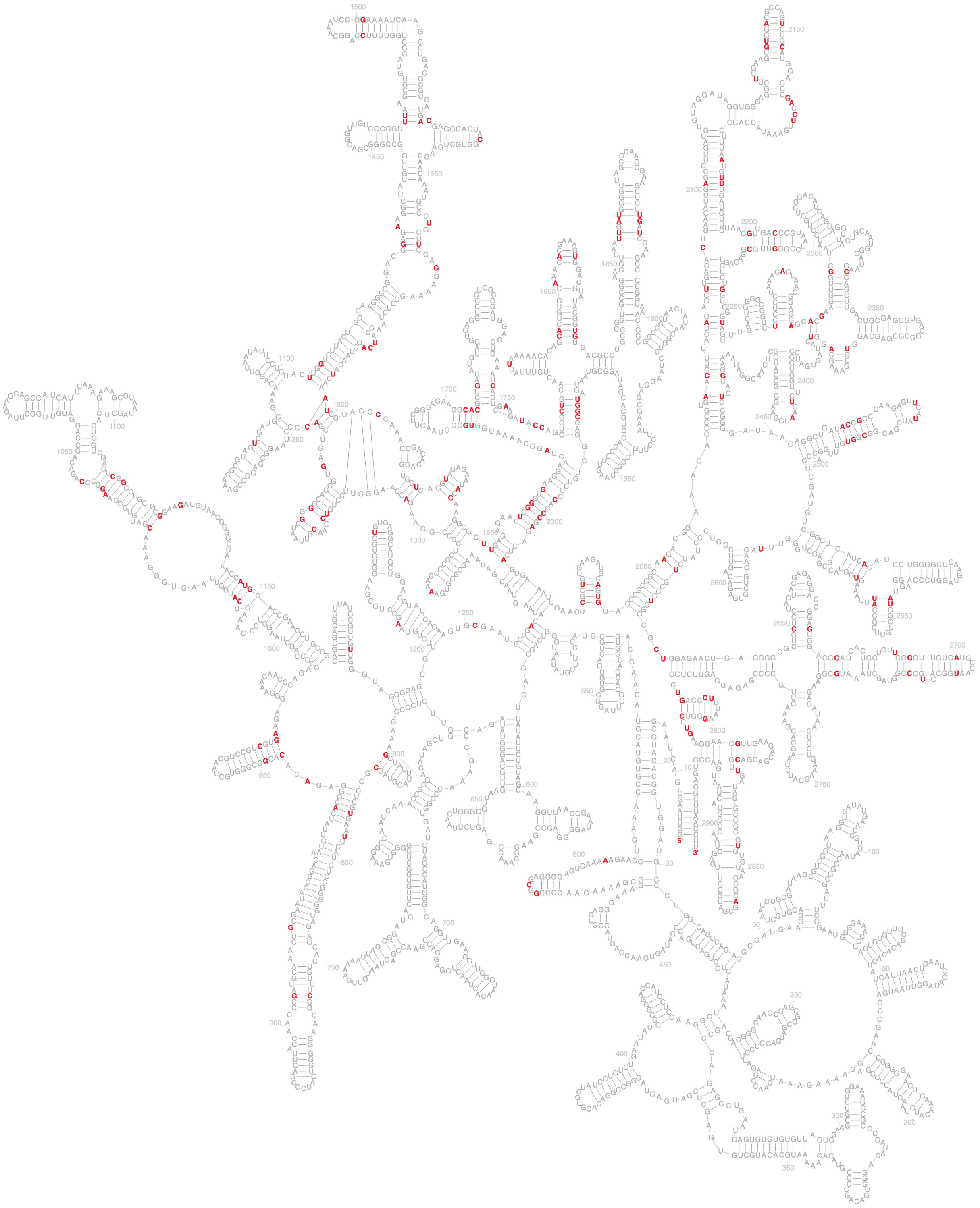

**Extended Data Fig. 9: Top 200 JSD locations from the 23S LM model.** Top ranking 200 Jensen-Shannon divergence values calculated from a secondary structure alignment of generated E. coli 23S rRNA sequences are colored in  red.

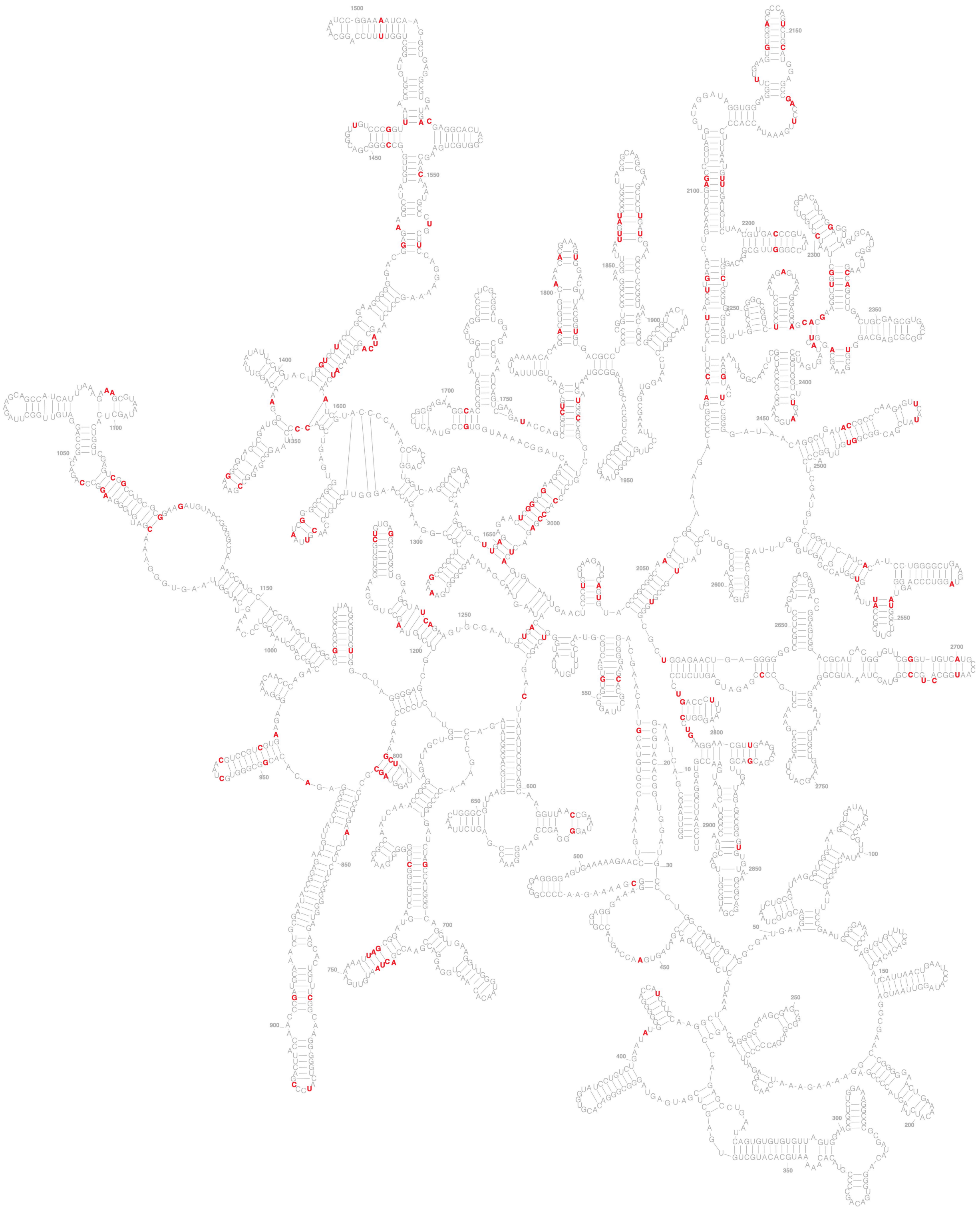

**Extended Data Fig. 10: Top 200 JSD locations from the 231 RNA LM model.** Top ranking 200 Jensen-Shannon divergence values calculated from a secondary structure alignment of generated E. coli 23S rRNA sequences are colored in red.

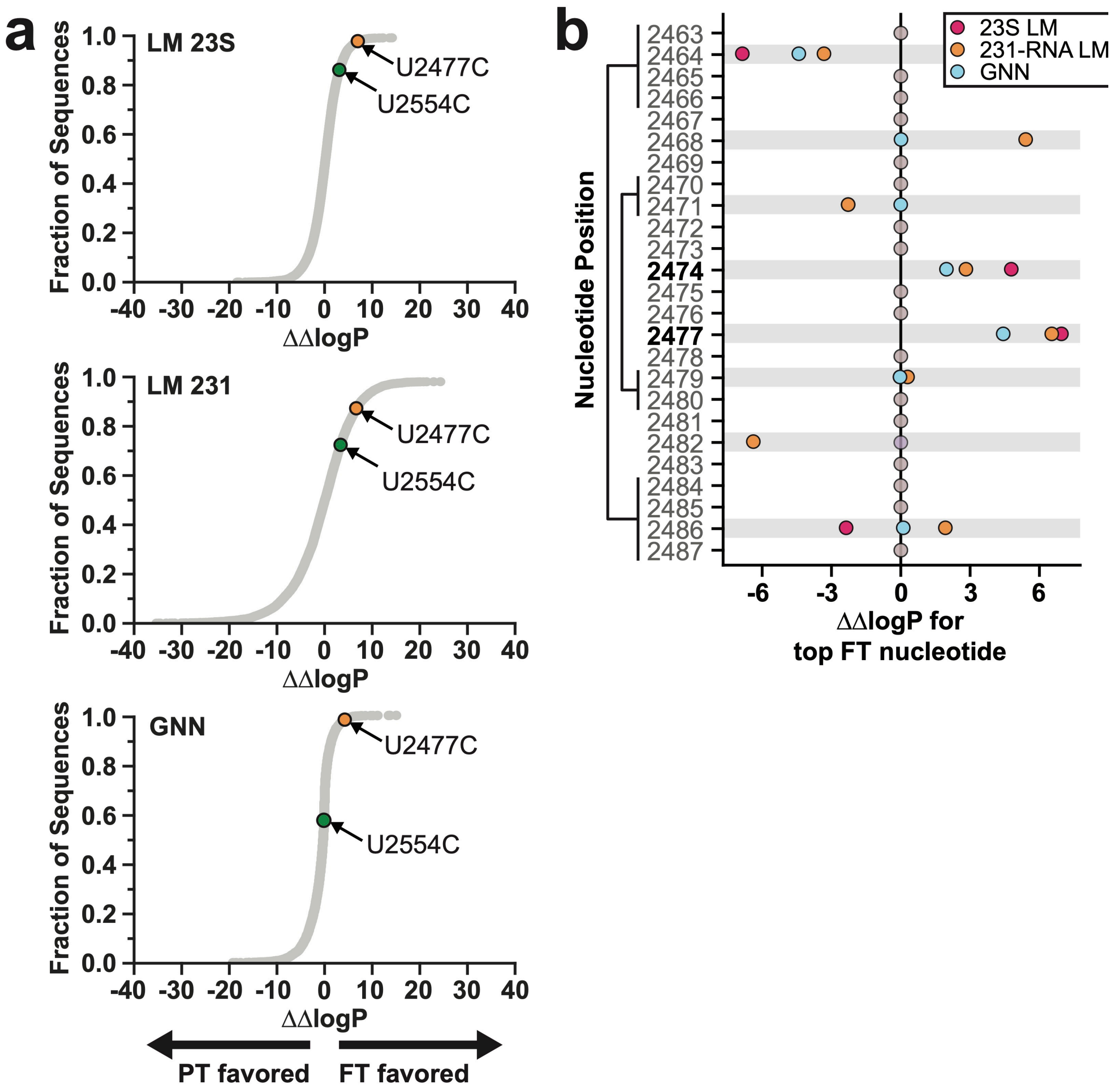


**Extended Data Fig. 11: Analysis of ΔΔlogP values for mutations in helix H89. a.** Cumulative plots of ΔΔlogP of all single mutations to the *E. coli* 23S rRNA for each model. Each point represents the probability of generating a single nucleotide mutant *E. coli* 23S sequence from the FT model relative to the PT model, normalized to that of the WT sequence. Two mutations, U2477C and U2554C, are denoted in orange and green, respectively. **b.** Analysis of helix 89 for candidate thermostabilizing mutations. For each position, the most frequent nucleotide in FT generated sequences (top FT nucleotide) is grafted into the *E. coli* 23S rRNA sequence and used to calculate ΔΔlogP(FT-PT) for the 23S LM, 231-RNA LM, and GNN models. Positions where the top FT nucleotide differs from WT in at least one model are highlighted in gray. Base pairing positions are indicated on the left.

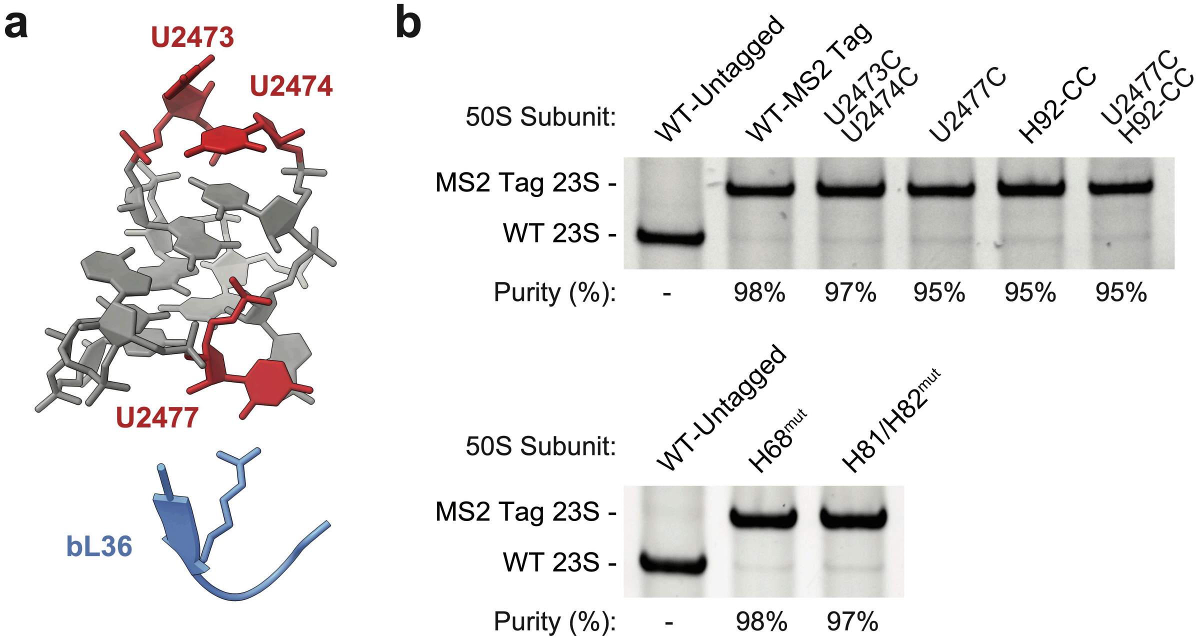


**Extended Data Fig. 12: Location and purification of rRNA mutations in the *E. coli* ribosome. a.** H89 and bL36 in the *E. coli* ribosome (PDB:7K00). 23S rRNA positions that are mutated in this study are shown in red. **b.** After 50S purification, 23S rRNA was isolated and subjected to RT-PCR analysis to quantify endogenous WT 50S contamination. The band intensities of RT-PCR products were used to quantify sample purity.

### Supplementary Information

**Supplementary Table 1: Information about the 231 RNA family dataset within GARNET.** The table includes Rfam model IDs, descriptions, and length, as well as the number of Rfam seed and full-alignment sequences. The table also includes the number of GARNET sequences for each Rfam ID, as well as the total GARNET sequence length.

**Supplementary Table 2: Optimal Growth Temperatures of GTDB organisms.** This dataset contains empirical OGTs from the TEMPURA and Gosha databases, OGTs predicted by TOME, GTDB genome accessions, GTDB taxonomic classifications, and thermal classifications for GTDB isolate descriptions.

**Supplementary Table 3: RNA language model parameters and training.** All pretrained language models used the following common hyperparameters: batch size = 18, dropout = 0.2, AdamW minimizer beta2 = 0.998, and use of the Flash attention algorithm[61]. Finetuned models had a batch size of 48. Extended parameters are also listed, including specifications for each GNN model.

**Supplementary Table 4: Jensen-Shannon divergences of RNA sequences generated by the RNA language models.** Nucleotide frequencies of natural sequences in GARNET and generated sequences are included, along with the JSD values at each position. *E. coli* 23S rRNA nucleotide numbering is shown, after removing gaps and insertions (**Materials and Methods**).

**Supplementary Table 5: Log likelihood calculations for candidate mutations in *E. coli* 23S rRNA.** Values were calculated using the strategy shown in **Fig. 6b**. Control calculations for all possible single-nucleotide mutations (GNN $\Delta\Delta$logP, 23S rRNA LM $\Delta\Delta$logP, and 231-RNA LM $\Delta\Delta$logP) are included.

**Supplementary Table 6: DNA sequences used in this study**. Comprehensive list of DNA sequences described in **Materials and Methods**.
